# CoCoA: Conditional Correlation Models with Association Size

**DOI:** 10.1101/2022.03.28.486098

**Authors:** Danni Tu, Bridget Mahony, Tyler M. Moore, Maxwell A. Bertolero, Aaron F. Alexander-Bloch, Ruben Gur, Dani S. Bassett, Theodore D. Satterthwaite, Armin Raznahan, Russell T. Shinohara

## Abstract

Many scientific questions can be formulated as hypotheses about conditional correlations. For instance, in tests of cognitive and physical performance, the trade-off between speed and accuracy motivates study of the two variables together. A natural question is whether speed-accuracy coupling depends on other variables, such as sustained attention. Classical regression techniques, which posit models in terms of covariates and outcomes, are insufficient to investigate the effect of a third variable on the symmetric relationship between speed and accuracy. In response, we propose CoCoA (Conditional Correlation Model with Association Size), a likelihood-based statistical framework to estimate the conditional correlation between speed and accuracy as a function of additional variables. We propose novel measures of the association size, which are analogous to effect sizes on the correlation scale, while adjusting for confound variables. In simulation studies, we compare likelihood-based estimators of conditional correlation to semi-parametric estimators adapted from genome association studies, and find that the former achieves lower bias and variance under both ideal settings and model assumption misspecification. Using neurocognitive data from the Philadelphia Neurodevelopmental Cohort, we demonstrate that greater sustained attention is associated with stronger speed-accuracy coupling in a complex reasoning task while controlling for age. By highlighting conditional correlations as the outcome of interest, our model provides complementary insights to traditional regression modelling and partitioned correlation analyses.

## 1. Introduction

When tasks depend on both speed and accuracy, neither measure alone can sufficiently capture performance and success (Gärtner and Strobel, 2021) due to the inherent need to prioritize either swiftness or correctness. Speed and accuracy are said to be coupled outcomes, and their mutual fluctuations result in speed-accuracy tradeoffs (SATs) (Heitz, 2014). The relationship between speed and accuracy is commonly thought to be monotonic and inversely related within individuals (Rahnev and Denison, 2018). However, individual tradeoff strategies can vary in study populations (for instance, in individuals with motor disorders (Fernani *and others*, 2017) or drug use disorders (de Dios *and others*, 2021)); SATs also change through development and aging (Forstmann *and others*, 2011; Mickeviciene *and others*, 2019), and can be modulated by task incentives (Manohar *and others*, 2015). Furthermore, speed and accuracy can become decoupled in cases when urgent decisions are needed (Salinas *and others*, 2014) or even positively related as a result of variations in attention (Bolsinova *and others*, 2016*a*).

To understand the drivers of speed-accuracy associations, psychometric studies are increasingly considering the two measures together (Boeck and Jeon, 2019). The simplest solution is to create a composite score using a (weighted) ratio or sum of speed and accuracy (Gur *and others*, 2014; Vandierendonck, 2021); however, combining them erases any fluid associations between speed and accuracy. Other sophisticated methods have considered the relationship hierarchically, with accuracy depending on response time (RT) and latent traits (Bolsinova *and others*, 2016*b*; Ranger and Ortner, 2012). While these models successfully represent the processes underlying RTs, they assess only an asymmetric relationship between speed and accuracy. For instance, consider that a linear model that regresses accuracy on speed and other covariates may give conflicting results compared to a model that regresses speed on accuracy and those same covariates.

Therefore, we conceptualize the speed-accuracy relationship in terms of the conditional correlation between variables, asking whether the strength and direction of this symmetric coupling depends on external factors, including attention and age. Unlike previous models that focus on a single unconditional correlation, we estimate the effect of covariates on this correlation (Loeys *and others*, 2011). To estimate model parameters, we identify three methods based on a Gaussian likelihood or a set of estimating equations, thereby avoiding the computational burden of Bayesian methods. In contrast to canonical correlation analyses (Parrington *and others*, 2014), a related method where the goal is to find maximally correlated linear combinations of variables, the target of our inference is the effect of a third variable (attention) on the correlation between speed and accuracy, after adjusting for other variables of interest or confounding variables (e.g., age).

Our proposed conditional correlation model with association size (CoCoA) adapts models from biostatistics (Wilding *and others*, 2011) and genome-wide association studies (Ho *and others*, 2010). We also propose novel measures of the effect size of attention on the speed-accuracy correlation, while adjusting for an arbitrary number of confounding variables. In simulations, we demonstrate that a simple restricted maximum-likelihood estimator is sufficient to achieve good results in both correctly and incorrectly specified data. This contradicts *a priori* expectations that more popular estimating equations-based methods would perform best, given their relative flexibility. Finally, we apply our framework to neurocognitive data from the Philadelphia Neurodevelopmental Cohort (Gur *and others*, 2012; Moore *and others*, 2015; Satterthwaite *and others*, 2016), demonstrating that increased attention tightens the coupling between speed and accuracy in a nonverbal reasoning task.

## 2. Conditional Correlation Models

### 2.1 Bivariate Gaussian Likelihood

Our main strategy is to consider bivariate distributions for RT and accuracy that contain mean, scale, and correlation parameters. Analogous to a regression model, where the mean parameter of that distribution would be modeled as a function of covariates, we also model the correlation and variance parameters as a function of covariates.

For each person *i*, we define their reaction time *X*_*i*_, accuracy *Y*_*i*_, attention *T*_*i*_, and age *Z*_*i*_. A natural choice for the distribution of **W**_*i*_ = (*X*_*i*_, *Y*_*i*_)*′* given *Z*_*i*_ and *T*_*i*_ is the bivariate conditional Gaussian distribution function:

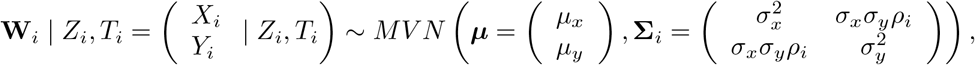

where the conditional correlation is modeled as

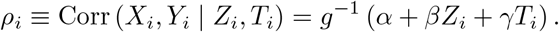

We choose the inverse link function *g*^−1^(·) to be the rescaled hyperbolic tangent function mapping real numbers to the [-1,1] interval, which allows unconstrained estimation of the conditional correlation parameters *α, β*, and *γ* (Bartlett, 1993):

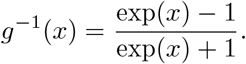

While the conditional correlation is the parameter of interest, we can also model the mean of the conditional bivariate distribution in terms of covariates:

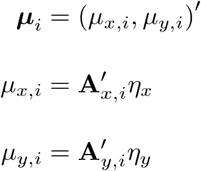

where **A**_*x,i*_ and **A**_*y,i*_ are covariate vectors for the *i*-th person, and *η*_*x*_ and *η*_*y*_ are their corresponding coefficients to be estimated. Similarly, we can also model the conditional variances

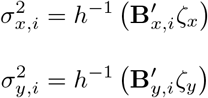

in terms of covariate vectors **B**_*x,i*_ and **B**_*y,i*_ and their coefficients *ζ*_*x*_ and *ζ*_*y*_. We choose the link function *h*^−1^(*x*) = log(*x*), which ensures that the variances are greater than 0. For bivariate distributions, the choice of *g*^−1^ and *h*^−1^ are sufficient to render the resulting 2 *×* 2 covariance matrix positive definite while allowing for unconstrained optimization of all model parameters.

Given observed data 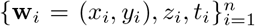 for *n* independent individuals, the overall log-likelihood is written as

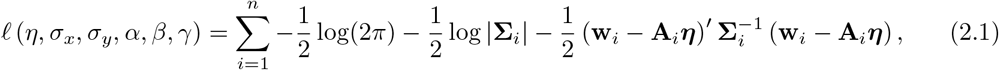

where ***η*** = (*η*_*x*_, *η*_*y*_)*′* and **A**_*i*_ is the design matrix for the bivariate mean vector defined by stacking 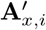 and 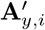 For simplicity, we assume that **B**_*x,i*_ = **B**_*y,i*_ = **1**; in principle, models for the conditional mean, variance, and correlation could depend on different or overlapping sets of covariates.

The maximum likelihood estimators (MLE) obtained from maximizing Equation (1) are known to be consistent but biased for the covariance, which is problematic for small samples. An unbiased estimate can be obtained from the restricted maximum likelihood (REML) estimator (Wilding et al., 2011), which maximizes a profile likelihood:

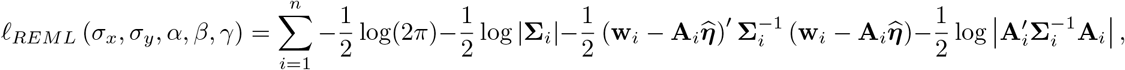

where 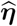 is the generalized least squares estimate of the mean parameters.

### 2.2 Estimating Equations

The bivariate distribution for (*X, Y* |*Z, T*) need not be fully specified to estimate conditional correlation parameters. A semi-parametric approach based on second-order estimating equations (GEE2) is often used in genome-wide association studies (Ho *and others*, 2010), where the goal is to assess how the correlation between expression levels of genes (co-expression) varies with cellular factors. The mean, variance, and correlation component models are the same as before:

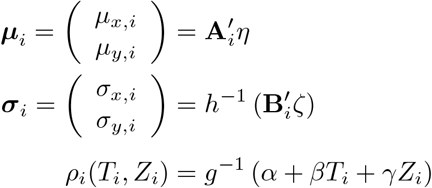

where **A**_*i*_ and **B**_*i*_ are design matrices for the mean and variance vectors. Importantly, the bivariate distribution of *X* and *Y* is not required to be Gaussian, and the parameters are not estimated by maximizing any likelihood function, but rather by solving a set of estimating equations (Yan and Fine, 2004).

### 2.3 Inference and Association Size

To determine if changes in attention *T* are significantly associated with changes in the conditional correlation between *X* and *Y*, we test the null hypothesis *H*_0_ : *γ* = 0. The estimate 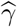 can be obtained by any of the 3 methods above (MLE, REML, or GEE2), and is valid whether *T* or other the covariates are binary or continuous. We estimate the sampling distribution of 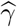 using a nonparametric bootstrap.

In practice, in addition to testing if the effect is significantly different from 0, interpretable effect sizes for these associations are critical. We refer to this quantity as the *association size*, because it describes the extent to which variations in *T* contribute to variations in the association between *X* and *Y*. While a natural candidate for association size would be 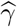 itself, this coefficient depends on the scale of *T* and is not simple to interpret given the nonlinear link function *g*^−1^.

When *T* is binary, we propose two measures of association size on the correlation scale, which will allow us to compare the contributions of different predictors. First, the marginal association size is defined as the expected correlation if *T* = 1 compared to *T* = 0, with the expectation taken over the marginal distribution of the confounding variable *Z*:

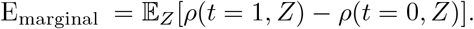

In other words, this is the expected change in correlation induced by varying *T* for the average person in our sample (Figure 1). Second, to account for the potential differences in the *T* = 0 and *T* = 1 groups, we also propose a conditional association size, defined as

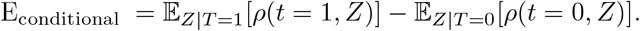

**Fig. 1.**
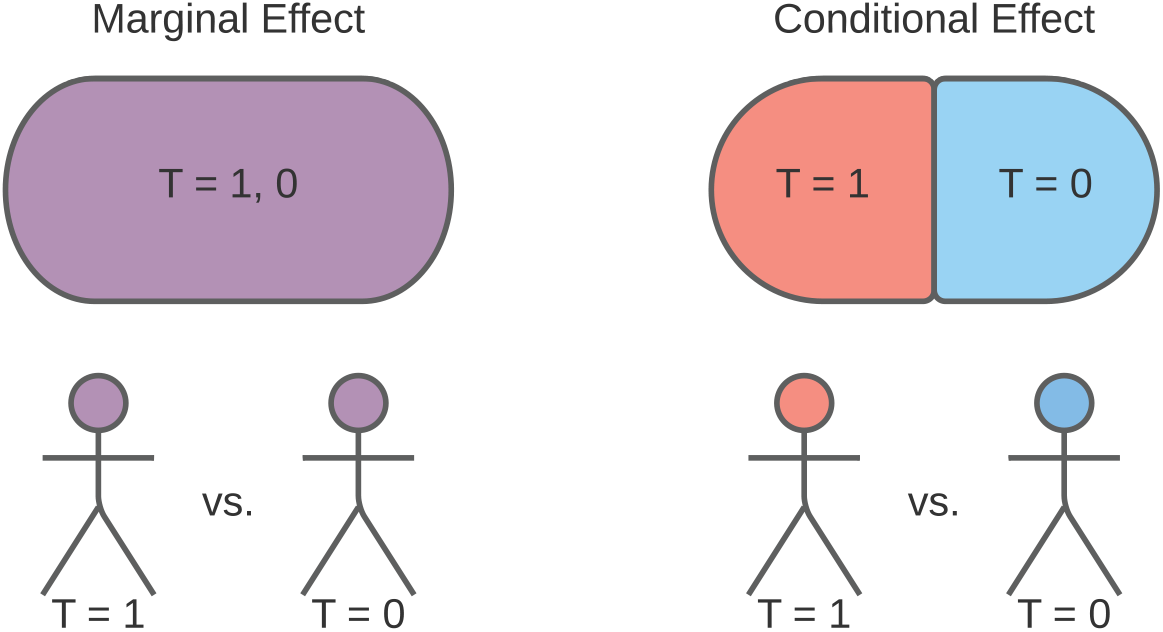
Marginal and conditional measures of association size. The marginal effect captures the change in conditional correlation between *X* and *Y* as a result of varying *T* in the study sample as a whole. The conditional correlation also considers potential confounding differences between the *T* = 0 and *T* = 1 subgroups induced by different conditional distributions of *Z* given *T*. When the average person in the *T* = 1 group is different from the average *T* = 0 group member (in terms of the confounding variable *Z*), the marginal effect and conditional effects will differ.

When the conditional distribution of *Z* differs f or *T* = 0 a nd *T* = 1, these contrasts will differ also. To estimate these effects, we use the plug-in estimator

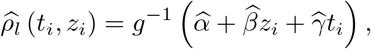

and replace the expectation with the sample mean

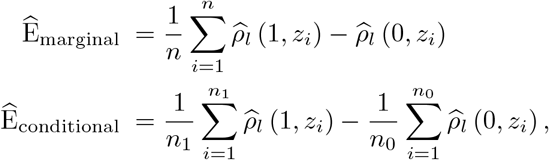

where *n*_1_ and *n*_0_ are the number of participants in the *T* = 1 and *T* = 0 groups, respectively. As before, confidence intervals for the effect sizes can be obtained by bootstrap methods.

### 2.4 Simulations

We compared the estimation accuracy of the MLE, REML, and GEE2 estimators in simulated data. All analyses were performed in R version 3.6.2 (R Core Team, 2019), with GEE2 estimation implemented in the *geepack* and *LiquidAssociation* packages. Within GEE2 estimators, we considered three types: that with full mean and scale component models 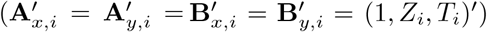; that with no mean component model (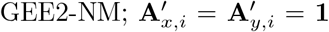 and 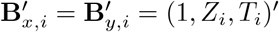), where all participants are constrained to have the same expected mean speed and accuracy; and that with no mean or scale component models (GEE2-NSM; 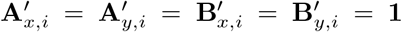), where all participants are constrained to have the same expected mean speed and accuracy and same variance components.

We drew samples of size *n* = 2000 and assumed, for convenience, that covariates were binary-valued:

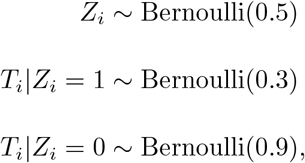

for *i* = 1, …, 2000. In the correctly specified condition, 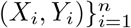 were then sampled from a bivariate normal distribution with means ***µ*** = (1, 1) and variances ***σ*** = (1, 2), and conditional correlation

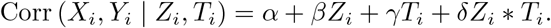

We allowed the parameter vector ***θ*** = (*α, β, γ, δ*) to take values in *{*0, 0.3, 0.6, 0.9, 1.2*} ×* **v** *×* **v** *×{*0, 0.3, 0, 7*}*, where **v** = *{*0, 0.075, 0.15, …, 1.425, 1.5*}*. For each unique point in the parameter space, this procedure resulted in one dataset on which we applied all estimators and computed the bias 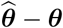.

We also considered the scenario in which the data-generating model was misspecified, i.e. when (*X*_*i*_, *Y*_*i*_) were drawn from a non-Gaussian distribution, but the MLE and REML estimators still relied upon a Gaussian likelihood. The misspecified distribution was defined using a copula distribution with Gamma(shape = 2, scale = 1) and Beta(shape = (1,1)) marginals, and the *Spearman* correlation defined as:

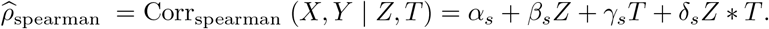

The subscript of the parameters ***θ***_*s*_ = (*α*_*s*_, *β*_*s*_, *γ*_*s*_, *δ*_*s*_) emphasize that the outcome here is the Spearman correlation, which is commonly used in defining copulas to denote dependence between two general random variables. We then transformed ***θ***_*s*_ to parameters corresponding to the Pearson correlation via the following procedure: given the estimate 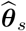, a sample of 100,000 points was sampled from the copula distribution defined above, using the Spearman correlation 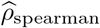 obtained by plugging in 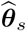 in the equation above. This procedure gives us the following system of equations:

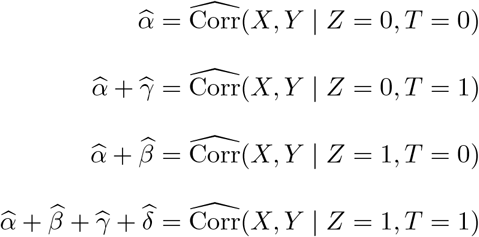

where 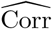 is the sample Pearson correlation. The solution to these equations 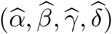 are the estimated parameters for the Pearson correlation.

## 3. Results

### 3.1 Simulation Results

In correctly specified simulated data, the MLE and REML estimators performed well compared to the three GEE2 estimators (Figure 2). Indeed, for many values of the parameter space, the GEE2 estimator did not converge, or the estimated parameter was more than several orders of magnitude larger than the true parameter. This was more likely to occur for larger values of the parameter space. Eliminating the mean model improved the convergence of the GEE2 estimator, but still resulted in larger bias compared to MLE and REML. While Figure 2 displays findings in a representative slice of the parameter space, we also calculated the performance across the entire parameter space by taking the median absolute difference 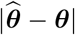, again finding that the REML estimation performed best (Table 1).

**Table 1.**
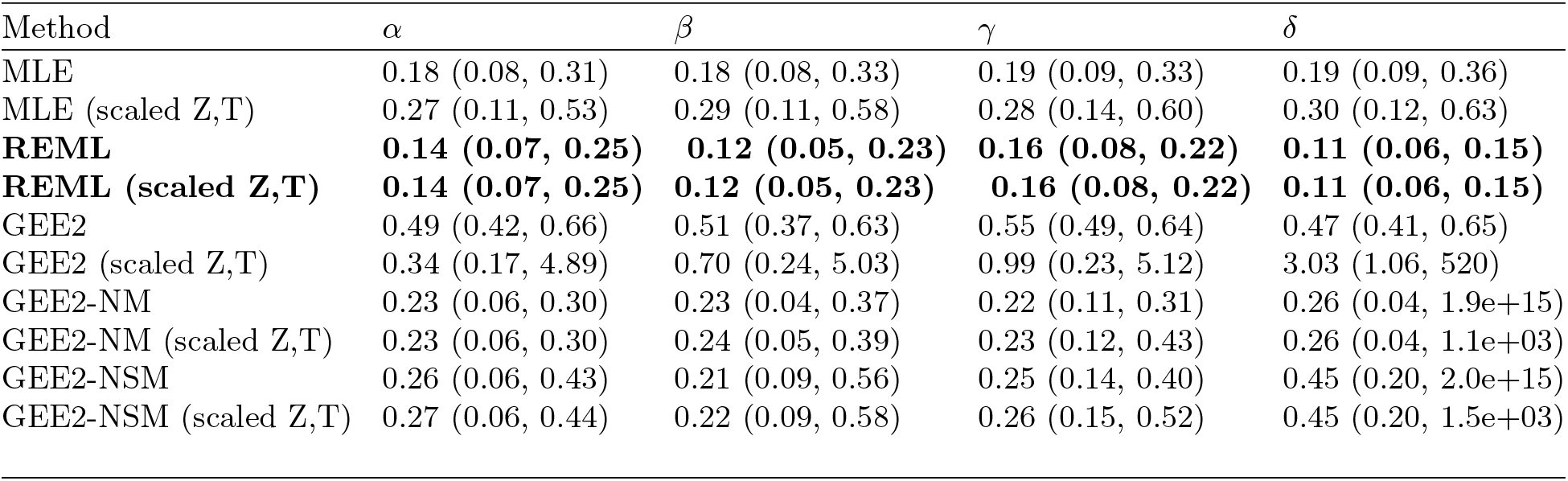
In simulated data, we assessed the performance of the MLE, REML, GEE2, GEE2-NM, and GEE2-NSM estimators using the absolute difference 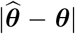 at all points in the parameter space. Each column corresponds to a parameter being estimated, and rows correspond to the estimator. Cell values are the median absolute difference (MAD), followed by the interquartile range (IQR) consisting of the 25th and 75th percentiles of absolute differences. GEE2 estimates were limited to those which converged. REML estimators performed best (bold cells), with lowest MAD and narrowest IQRs, and were not substantially affected by z-scoring the covariates Z and T. (Performance was identical up to 4 decimal places.) However, the MLE and all GEE2 estimators were affected by z-scoring, with the latter displaying greater variability in MAD and IQR.

**Fig. 2.**
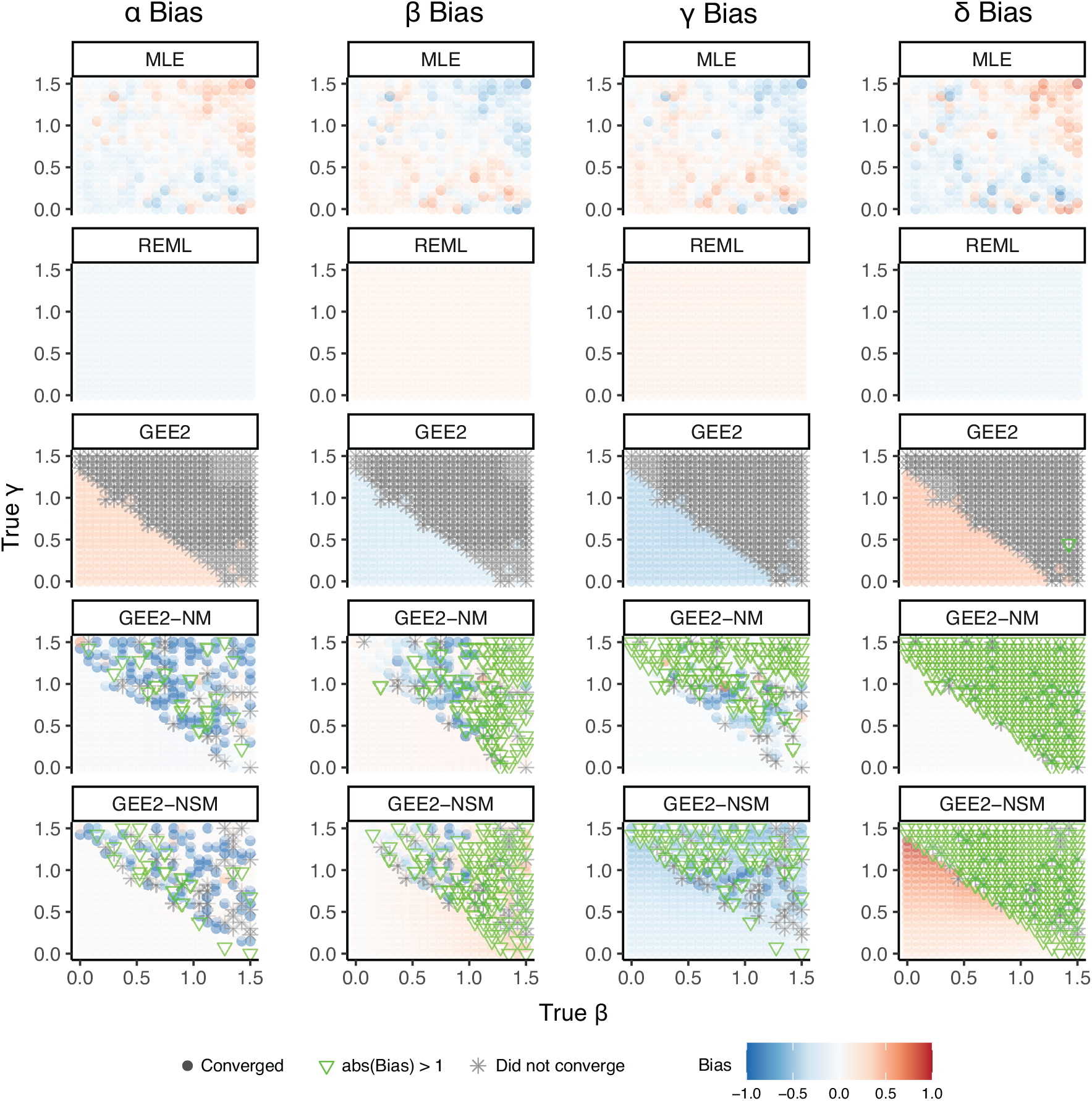
In simulated data, the REML and MLE estimators perform better for all model parameters, compared to the GEE2 estimators. In these bias plots, the true value of (*α, δ*) were fixed at (0.9,0.3), while the true values of (*β, γ*) varied along the *x*- and *y*-axes. Each column corresponds to each of the 4 model parameters, and each row corresponds to a different estimator. For a given point in the parameter space (*α*_0_, *β*_0_, *γ*_0_, *δ*_0_), a single simulated dataset of 2000 points was generated using those parameters, and all estimators were calculated on that same data. The color of the dot represents the bias (the difference between the estimated and true value) of that parameter estimate. We considered bias values between −1 and 1 to highlight meaningful differences between estimators; green hollow triangles represent estimates that exceeded an absolute bias of 1, and gray asterisks signify cases where the estimator did not converge. Overall, we found that the REML estimator performed best in minimizing error, while the full GEE2 estimator was often affected by convergence issues. The GEE2-NM and GEE2-NSM estimators, derived by excluding either the mean model or the mean and scale models, somewhat ameliorated these convergence issues.

In theory, the process of *z*-scoring (i.e, linearly scaling by the mean and standard deviation) of the covariates *Z* and *T* should not affect the overall parameter estimation after back-transforming to the original scale. While we observed no substantial impact of *z*-scoring on REML estimation, both MLE and GEE2 estimation were differentially impacted: GEE2 estimation improved overall, while MLE estimation worsened (Table 1). Finally, for all models, we also examined the interquartile range (IQR) of absolute differences 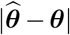, defined by the 25th and 75th percentile. Overall, the REML estimator had the narrowest IQR, while the IQR of the GEE2 estimators proved to be orders of magnitude larger, signifying that the estimator may have suffered from high variability for a large portion of the parameter space.

When the data-generating model was misspecified, the REML estimator again performed best on average (Table S1), although we expected GEE2 to perform best under non-Gaussian data. However, the IQRs for GEE2 estimates were more narrow compared to those in the Gaussian data. For the slice of the parameter space where (*α, δ*) = (0.9, 0.3), the GEE2 estimators converged more often (Figure S1) when the model was misspecified.

### 3.2 Speed/Accuracy Trade-offs in Complex Reasoning Tasks

We next applied our method to data from the Philadelphia Neurodevelopmental Cohort (Gur *and others*, 2012; Satterthwaite *and others*, 2016), a large-scale study of brain function and behavior in adolescents. Specifically, we investigated the coupling between speed and accuracy in individuals taking the Penn Matrix Reasoning Test (PMRT), which is designed to assess nonverbal reasoning as part of the Penn Computerized Neurocognitive Battery (Gur *and others*, 2010). In total, *n* = 3879 participants answered up to 24 items each. For each participant, their overall accuracy was calculated as the proportion of correct responses, and speed was represented by the log-transformed median RT. We used the median to be consistent with prior studies (Moore *and others*, 2015), though the distribution of the log-transformed mean RT was similar. The log-transform was used because it resulted in a less skewed distribution of RTs, consistent with the common assumption that RTs follow a log-normal distribution (van der Linden, 2009), and the resulting response times had a linear relationship with accuracy (Figure 3).

**Fig. 3.**
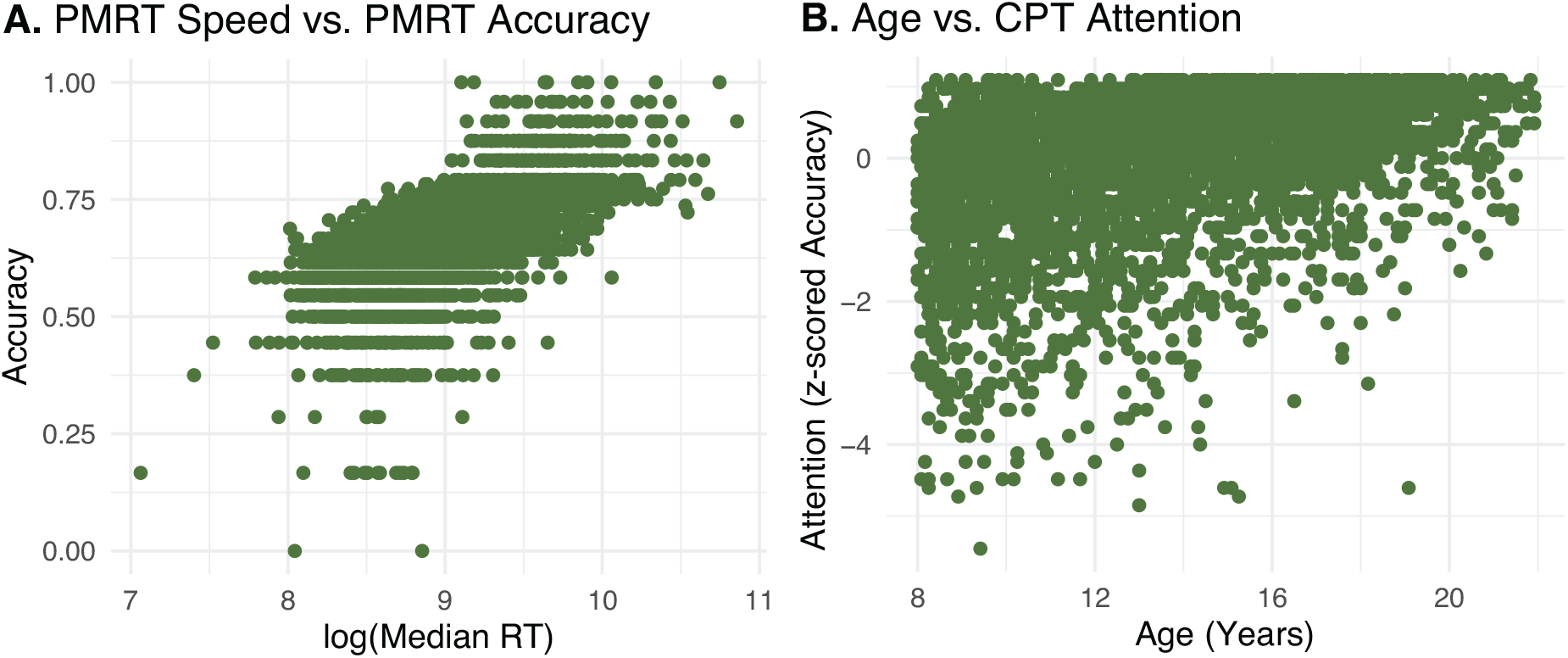
Scatterplots of performance and individual traits in neurocognitive data from the Philadelphia Neurodevelopmental Cohort (*n* = 3879). Panel A: the relationship between speed (log-transformed RT) and accuracy from the Penn Matrix Reasoning Task (Version A) is approximately linear and positive, reflecting a trade-off between swiftness and correctness. Panel B: attention was assessed using an accuracy measure on the Continuous Performance Test, which assesses vigilant attention. Age is differentially distributed in lower and higher attention groups, and is included in the conditional correlation model as a confound variable.

In addition to speed and accuracy on the PMRT, attention was assessed using the Continuous Performance Test (CPT), a task requiring vigilant attention that presents participants with 7-segment displays at 1 Hertz and instructs them to press the space bar when the display forms either a number (Numbers condition) or letter (Letters condition). The CPT is not administered simultaneously to the PMRT, but typically takes place within the same 1-hour testing session. We used a participant’s *z*-scored CPT accuracy as a proxy for attention, and binarized the sample into top (high-attention) and bottom (low-attention) tertiles. This produced a final sample of size *n* = 2880 individuals. Finally, we included age as a confounding variable in our model, since it is differentially distributed in low- and high-attention groups, and is also associated with the SAT (Figure 3).

Letting *X* be the log-transformed median RT on the PMRT, *Y* be the proportion of accurate responses on the PMRT, *Z* be the age in years, and *T* be the high or low attention group, the final conditional correlation model was

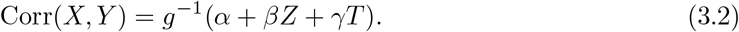

We considered MLE, REML, GEE2-NM, and GEE2-NSM estimates of the model parameters and the association sizes defined in the previous section. Due to excessive variability, the full GEE2 estimates were not included in this analysis. As hypothesized, we found that greater attention corresponded to stronger speed-accuracy coupling, a small but significant increase (Figure 4). Among the MLE and REML estimators, nearly all marginal effects and conditional effects were significantly greater than 0 and centered around 0.12, with conditional effects being slightly greater. The GEE2-NSM estimates of effect had larger variability, likely due to increased variability in estimating model parameters (Figure S2). Both GEE2 estimates were closer to 0 or centered around −0.1, in contrast to the other estimators and sliding window correlation, a nonparametric measure of conditional correlation (Figure S3).

**Fig. 4.**
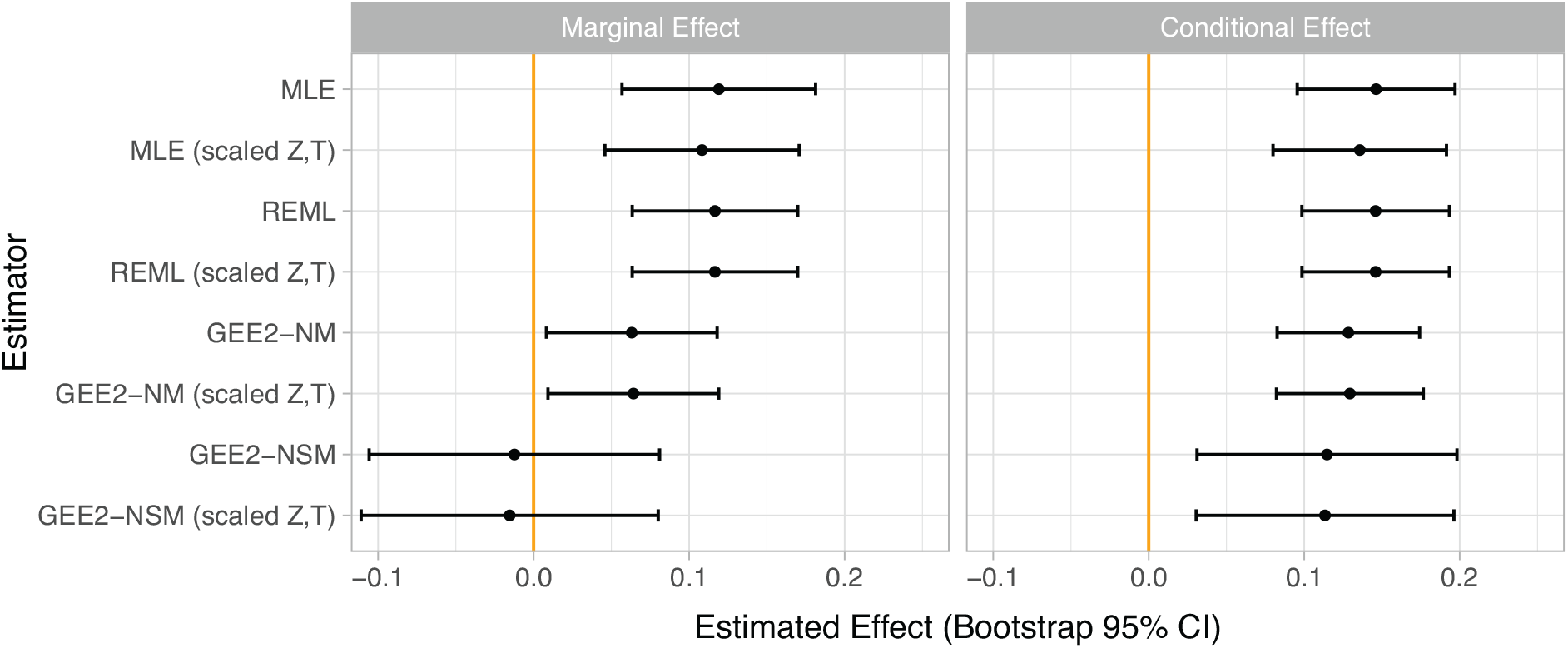
Estimated effects of high versus low attention, defined by the top and bottom tertiles of attention score, on speed-accuracy coupling in the PMRT (Version A). Using the MLE and REML estimators, a stronger speed-accuracy coupling was estimated for higher vs. lower attention groups. However, we found more variable or null results from the GEE2-NM and GEE2-NSM estimators, particularly in estimating the marginal effect.

## 4. Discussion

We introduced and validated a framework for modelling conditional correlations as a function of multiple covariates, and proposed a novel association size that adjusts for confounding. This proposed method allows us to assess the symmetric coupling between two outcomes of interest, providing complementary insights to analyses that model one outcome as a function of the other. In comparison to partitioned correlation analyses or nonparametric methods such as sliding correlation, our regression-based framework has the advantage that the coupled outcome can depend on an arbitrary number of both continuous and categorical variables. While the association size was only defined for binary *T*, continuous extensions could follow in the vein of dose-response curves or correlations commonly used as effect sizes for continuous treatments (Rutledge and Loh, 2004).

We identified three candidate methods for model estimation. In simulation studies, we found that the likelihood-based estimators (MLE, REML) were more accurate and less variable compared to the semiparametric estimators (GEE2). Between the full GEE2 model and those without mean component models (GEE2-NM and GEE2-NSM), we observed that removing the mean model often improved estimation bias and numerical convergence, suggesting that the full model may be overspecified. We also conjecture that numerical instability could result from the choice of the link function *g*^−1^(*x*), due to the flatter shape of the function at lower and higher extremes. Under a misspecified model condition, these findings persisted, although with the caveat that model was only misspecified in one particular way. However, our findings show that distributional assumptions do not necessarily have to be met in order for MLE and REML to perform well.

Next, we used our model to ask whether the coupling between speed and accuracy depends on attention. Given the MLE and REML estimators, we concluded that higher attention groups indeed have stronger coupling, corroborating nonparametric measures of conditional correlation as well as previous findings in hierarchical models (Meng *and others*, 2015). However, our findings using the GEE2, GEE2-NM, and GEE2-NSM estimators were more variable or closer to the null. Together with the simulation results, and other rare reports of high variability in GEE2 estimates (Chen *and others*, 2020; Franke *and others*, 2004), we recommend excluding the mean model if these parameters are not important to the scientific inquiry and if estimates are overly large. We also recommend, whenever possible, to form conclusions based on an aggregate of different methods.

The relationship between speed and accuracy is nuanced and is a topic of study in its own right. Our model did not account for differences between low-RT (“easy”) and high-RT (“difficult”) questions, which are thought to correspond to different processes (Bolsinova *and others*, 2016*a*). In the context of SAT, it is important to distinguish between within-individual and inter-individual correlations, since they are often reversed (Boeck and Jeon, 2019): speed and accuracy often have positive correlation cross-sectionally but negative correlation within individuals or groups, a phenomenon known as Simpson’s paradox (Kievit *and others*, 2013). Because we aggregated accuracy and reaction time data for each individual, our method is only able to measure interindividual correlations, though we look forward to future work that can accommodate correlated observations within individuals.

## 5. Software

Software in the form of R code, together with documentation and instructions on replicating all Tables/Figures, are available on request from the corresponding author, and will be made publicly available on GitHub upon publication.

## 6. Supplementary Material

Supplementary material is available online at http://biostatistics.oxfordjournals.org.

## Acknowledgments

This work was supported by National Institutes of Health [grant numbers R01MH112847 to R.T.S. and T.D.S.; and R01MH120482, R01EB022573, and R37MH125829 to to T.D.S.]

