## Supplemental Figures and Tables for "CoCoA: Conditional Correlation Models with Association Size"

Supplementary Material to
"CoCoA: Conditional Correlation Models with Association Size"

Danni Tu[1], Bridget Mahony[2], Tyler M. Moore[3], Maxwell A. Bertolero[3,4], Aaron F. Alexander-Bloch[3], Ruben Gur[3], Dani S. Bassett[5,6,7,8], Theodore D. Satterthwaite[3,4], Armin Raznahan[2], Russell T. Shinohara[1]

[1] Penn Statistics in Imaging and Visualization Center, Department of Biostatistics, Epidemiology, and Informatics, Perelman School of Medicine, University of Pennsylvania, Philadelphia, PA, 19104
[2] Section on Developmental Neurogenomics, National Institutes of Mental Health, Bethesda, MD, 20814
[3] Department of Psychiatry, Perelman School of Medicine, Philadelphia, PA, 19104
[4] Penn Lifespan Informatics and Neuroimaging Center, Philadelphia, PA, 19104
[5] Department of Bioengineering, University of Pennsylvania, Philadelphia, PA, 19104
[6] Department of Physics and Astronomy, University of Pennsylvania, Philadelphia, PA, 19104
[7] Department of Electrical and Systems Engineering, University of Pennsylvania, Philadelphia, PA, 19104
[8] Department of Neurology, University of Pennsylvania, Philadelphia, PA, 19104

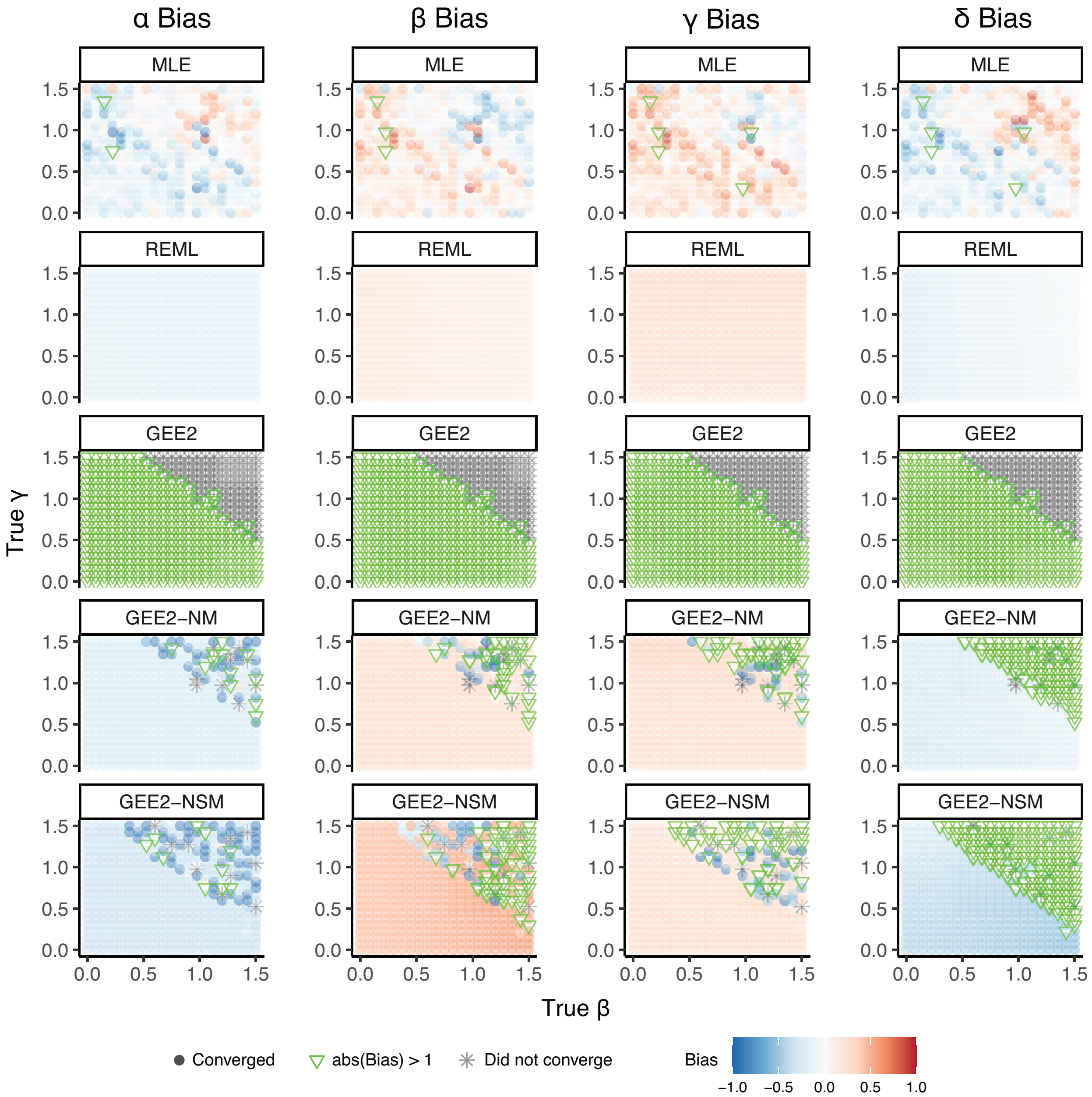


Figure S1. In an instance where the bivariate distribution of $(X, Y|Z, T)$ was misspecified, REML and MLE estimators continued to perform well. The GEE2-NM and GEE2-NSM estimators also performed comparably, with occasionally high variability, while the full GEE2 estimator performed less well compared to the Gaussian data. These bias plots represent a slice of the parameter space, with the true value of (α,δ) fixed at (0.9,0.3), while the true values of (β,γ) varied along the x- and y-axes. Each column corresponds to each of the 4 model parameters, and each row corresponds to a different estimator. For a given point in the parameter space $(\alpha_0, \beta_0, \gamma_0, \delta_0)$, a single simulated dataset of 2000 points was generated using these parameters, and all estimators were calculated on the same data. The color of the dot represents the bias (the difference between the estimated and true value) of the parameter estimate. We considered bias values between -1 and 1 to highlight meaningful differences between estimators; green hollow triangles represent estimates that exceeded an absolute bias of 1, and gray asterisks signify cases where the estimator did not converge.

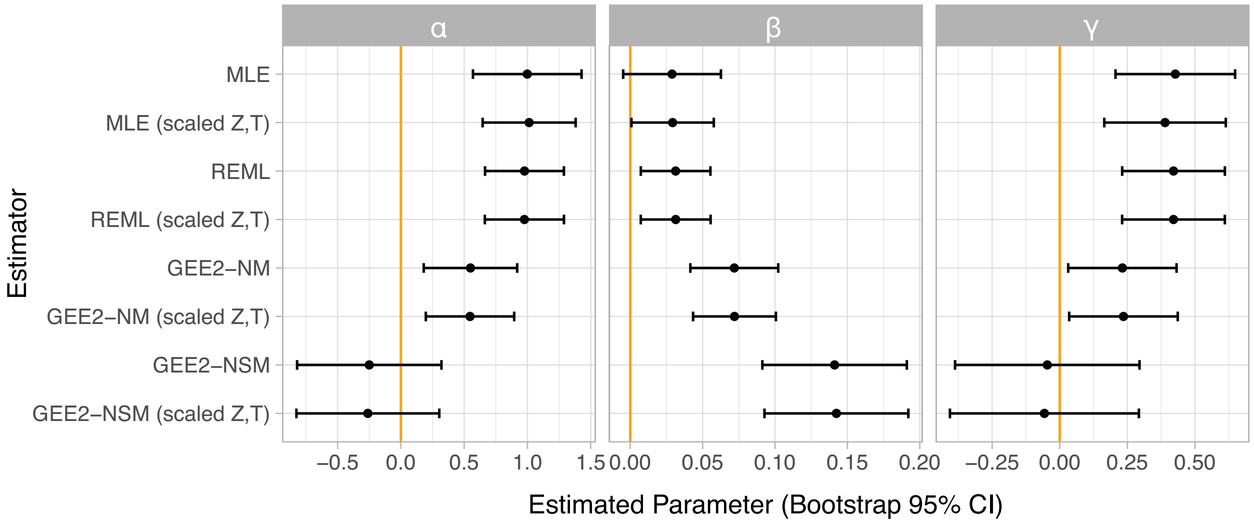


Figure S2. Estimated model parameters of the conditional correlation model in Equation (2). Much of the variability of the association sizes when utilizing the GEE2-NM and GEE-NSM estimators (Figure 4) can be attributed to variability in estimating $\alpha$ and $\gamma$, with the latter driving discrepancies in the sign of the marginal association size.

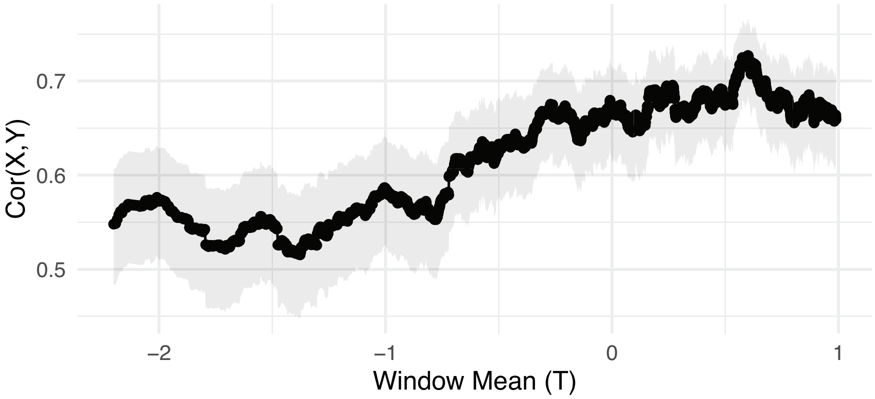


Figure S3. The conditional correlation between *X* (log-transformed median RT) and *Y* (accuracy) on the PMRT Version A as a function of sliding windows of *T* (attention), a nonparametric measure of conditional association. In individuals corresponding to the top tertile of attention, speed-accuracy coupling is higher by around 0.1.

| **Method** | $\boldsymbol{\alpha}$ | $\boldsymbol{\beta}$ | $\boldsymbol{\gamma}$ | $\boldsymbol{\delta}$ |
|---|---|---|---|---|
| MLE | 0.16<br>(0.07, 0.29) | 0.17<br>(0.08, 0.30) | 0.19<br>(0.09, 0.33) | 0.23<br>(0.11, 0.38) |
| MLE (scaled Z,T) | 0.22<br>(0.10, 0.42) | 0.25<br>(0.11, 0.47) | 0.28<br>(0.13, 0.49) | 0.31<br>(0.14, 0.56) |
| **REML** | **0.11**<br>**(0.04, 0.21)** | **0.14**<br>**(0.04, 0.22)** | **0.15**<br>**(0.09, 0.33)** | **0.20**<br>**(0.10, 0.38)** |
| **REML (scaled Z,T)** | **0.11**<br>**(0.04, 0.21)** | **0.14**<br>**(0.04, 0.22)** | **0.15**<br>**(0.09, 0.33)** | **0.20**<br>**(0.10, 0.38)** |
| GEE2 | 1.69<br>(1.56, 1.86) | 1.73<br>(1.63, 1.87) | 1.72<br>(1.48, 1.90) | 1.73<br>(1.51, 1.90) |
| GEE2 (scaled Z,T) | 0.16<br>(0.06, 0.24) | 0.26<br>(0.07, 0.34) | 0.19<br>(0.09, 0.35) | 4.58<br>(4.20, 4.87) |
| GEE2-NM | 0.19<br>(0.07, 0.27) | 0.21<br>(0.06, 0.30) | 0.20<br>(0.09, 0.35) | 0.25<br>(0.12, 0.33) |
| GEE2-NM (scaled Z,T) | 0.19<br>(0.07, 0.29) | 0.23<br>(0.06, 0.30) | 0.20<br>(0.09, 0.36) | 0.25<br>(0.12, 0.33) |
| GEE2-NSM | 0.20<br>(0.07, 0.36) | 0.16<br>(0.05, 0.43) | 0.26<br>(0.12, 0.43) | 0.40<br>(0.19, 0.91) |
| GEE2-NSM (scaled Z,T) | 0.20<br>(0.07, 0.38) | 0.17<br>(0.05, 0.45) | 0.27<br>(0.12, 0.50) | 0.40<br>(0.19, 0.88) |

Table S1. Under the misspecified model, we assessed the performance of the MLE, REML, GEE2, GEE2-NM, and GEE-NSM estimators using the absolute difference $|\hat{\theta} - \theta|$ at all points in the parameter space. GEE2 estimates were limited to those which converged. Each column corresponds to a parameter being estimated, and rows correspond to the estimator. Cell values are the median absolute difference (MAD), followed by the interquartile range (IQR) consisting of 25th and 75th percentiles of absolute differences. As in the correctly specified case, REML estimators performed best (bold cells), with lowest MAD and narrowest IQRs. REML estimation was not substantially affected by *z*-scoring the covariates *Z* and *T*. (Performance was identical up to 4 decimal places.) The GEE2, GEE2-NM, and GEE-NSM estimators appeared to perform better in the misspecified case compared to in the Gaussian data, exhibiting smaller IQRs.
